# Epitope-resolved profiling of the SARS-CoV-2 antibody response identifies cross-reactivity with an endemic human CoV

**DOI:** 10.1101/2020.07.27.222943

**Authors:** Jason T Ladner, Sierra N Henson, Annalee S Boyle, Anna L Engelbrektson, Zane W Fink, Fatima Rahee, Jonathan D’ambrozio, Kurt E Schaecher, Mars Stone, Wenjuan Dong, Sanjeet Dadwal, Jianhua Yu, Michael A Caligiuri, Piotr Cieplak, Magnar Bjørås, Mona H Fenstad, Svein A Nordbø, Denis E Kainov, Norihito Muranaka, Mark S Chee, Sergey A Shiryaev, John A Altin

## Abstract

A high-resolution understanding of the antibody response to SARS-CoV-2 is important for the design of effective diagnostics, vaccines and therapeutics. However, SARS-CoV-2 antibody epitopes remain largely uncharacterized, and it is unknown whether and how the response may cross-react with related viruses. Here, we use a multiplexed peptide assay (‘PepSeq’) to generate an epitope-resolved view of reactivity across all human coronaviruses. PepSeq accurately detects SARS-CoV-2 exposure and resolves epitopes across the Spike and Nucleocapsid proteins. Two of these represent recurrent reactivities to conserved, functionally-important sites in the Spike S2 subunit, regions that we show are also targeted for the endemic coronaviruses in pre-pandemic controls. At one of these sites, we demonstrate that the SARS-CoV-2 response strongly and recurrently cross-reacts with the endemic virus hCoV-OC43. Our analyses reveal new diagnostic and therapeutic targets, including a site at which SARS-CoV-2 may recruit common pre-existing antibodies and with the potential for broadly-neutralizing responses.

## Introduction

SARS-CoV-2 is a single-stranded RNA virus in the Coronaviridae family that emerged in late 2019 and has caused morbidity, mortality and economic disruption on a global scale with few precedents (Zhu et al., 2020). The Coronaviridae family includes four species/strains that are endemic in the human population and usually associated with mild, self-limiting upper respiratory tract infections: HCoV-229E, HCoV-NL63, HCoV-HKU1 and HCoV-OC43 (Betacoronavirus 1 species). Two other species – MERS-CoV and SARS-CoV– have recently emerged to cause severe disease in humans. Like the other human-infecting coronaviruses (CoV) (Callow et al., 1990; Dijkman et al., 2008), SARS-CoV-2 infection can elicit a robust antibody response in humans (Liu et al., 2020; Ni et al., 2020) and this response represents the major focus of widespread efforts to develop accurate diagnostics, as well as strategies for passive and active immunization against infection (Casadevall and Pirofski, 2020; Krammer and Simon, 2020; Thanh Le et al., 2020). Existing serological assays for SARS-CoV-2 antibody reactivity generally use full-length viral proteins or domains – Spike (S), Nucleocapsid (N), or the receptor-binding domain (RBD) of S – as antigenic baits, followed by enzyme-linked or fluorescent detection (Krammer and Simon, 2020). These assays provide a single measure of antibody reactivity, which represents a composite signal across many epitopes, and are able to detect viral exposure with a range of accuracies (Deeks et al., 2020; Whitman et al., 2020). Neutralization assays using either native or pseudotyped viruses have also been developed (Nie et al., 2020). It remains to be seen how these different assays will perform as diagnostics or correlates of the protection conferred by infection or vaccination.

Relative to protein-based analyses of the humoral response, epitope-level assays have the potential to add several layers of information. First, although SARS-CoV-2 proteins are generally distinct from other human-infecting Coronaviruses, some regions of strong homology exist (Lu et al., 2020; Zhu et al., 2020), meaning that there is the potential for immune cross-reactivity that can only be resolved at the epitope level. Indeed, it was recently demonstrated that a large fraction of non-exposed individuals have T cell reactivity to SARS-CoV-2 peptides, indicating cross-reactivity with existing responses, possibly those generated against homologous peptides from endemic CoVs (Grifoni et al., 2020). In the case of antibody responses, cross-reactivity has been described between the more closely related SARS-CoV and SARS-CoV-2 (Lv et al., 2020; Pinto et al., 2020). Epitope-resolved analyses therefore have the potential to identify antigens that may discriminate related CoVs, leading to more specific diagnostic assays. High levels of sequence conservation may also indicate functional essentiality; therefore, by highlighting potentially cross-reactive epitopes in conserved regions of the proteome, epitope-level assays can identify antibodies and targets with therapeutic potential, against which viral escape may be more difficult (Friesen et al., 2014).

A second rationale for generating epitope-resolved views is that antibody recognition of different protein regions can have divergent functional consequences, including neutralization potential. For coronaviruses, antibodies binding the surface-exposed, receptor-binding S protein exhibit the greatest neutralizing potential (Du et al., 2009; Pillay, 2020), but these antibodies can recognize a wide variety of epitopes within the protein, each with the potential for different functional consequences. This likely accounts for the imperfect correlation between the titers of S-binding antibodies and viral neutralization activity across individuals (Robbiani et al., 2020). Due to its interaction with the host entry receptor (the angiotensin converting enzyme 2 – ACE2), the RBD of S represents the predominant target of vaccination and monoclonal antibody development strategies, and a growing number of antibodies against this domain have been described (Chi et al., 2020; Hansen et al., 2020; Robbiani et al., 2020; Zost et al., 2020). However, the RBD is one of the less conserved regions of the CoV proteome and antibodies against epitopes outside of the RBD have also been shown to have neutralizing activity (Chi et al., 2020; Poh et al., 2020): these may act in various ways, including by preventing important protease cleavage events and/or conformational changes required for successful entry into cells. On the other hand, antibodies that recognize epitopes within the N protein, which coats the viral genome and is contained within mature viral particles, likely provide little or no neutralization potential, but may be useful signatures for differentiating vaccine responses from those resulting from natural virus infection, a strategy already used for other viruses (Hoofnagle et al., 1974; Lubroth et al., 1996). In addition to different neutralization potential, it is possible that unfavorable distributions of epitope reactivity can contribute to immunopathology, for example through antibody dependent enhancement (Halstead and O’rourke, 1977; Katzelnick et al., 2017; Khurana et al., 2013), although this phenomenon remains to be demonstrated for SARS-CoV-2 (Eroshenko et al., 2020).

Peptide sub-sequences have been used for decades as probes to detect antibodies recognizing linear epitopes within the full-length proteins from which they are derived (Fleri et al., 2017; Lucchese et al., 2007). Although unable to detect antibodies whose binding depends on elements that are discontinuous in the primary sequence, this strategy has the advantage that it enables the highly-efficient design and synthesis of antigen baits. In its simplest format, peptides can be used individually, for example in separate wells in an ELISA. A recent study used this approach to identify two linear epitopes in S protein that were targeted by neutralizing antibodies in SARS-CoV-2 convalescent donors (Poh et al., 2020). More powerful assays involve sets of peptides that are assayed in multiplex – using either spatial addressing, in the case of peptide arrays (Price et al., 2012), or DNA indexing, in the case of phage display libraries (Larman et al., 2011). Using the latter approach, the highly-multiplexed and epitope-resolved detection of antibodies to viruses has been demonstrated with high sensitivity and specificity (Xu et al., 2015).

Here, we present a new, synthetic biology approach to highly-multiplexed peptide-based serological assays (‘PepSeq’) in which libraries of peptide baits – each covalently coupled to a DNA barcode – are synthesized from high-complexity DNA pools using a simple and fully *in vitro* approach. Library synthesis takes advantage of *in vitro* transcription and translation, including an intramolecular coupling mediated by puromycin (Kozlov et al., 2012; Roberts and Szostak, 1997) and the DNA-barcoded peptides can then be used to probe antibodies using a high-throughput sequencing read-out. We use this platform to synthesize libraries of overlapping 30mer peptides covering all human CoV proteomes and assay these against sera from pre-pandemic and SARS-CoV-2 convalescent donors. Our results demonstrate accurate detection of SARS-CoV-2 exposure and reveal multiple immunodominant antibody epitopes, including at least one in which antibody responses cross-react between SARS-CoV-2 and an endemic human CoV.

## Results

### A highly-multiplexed peptide assay to evaluate CoV antibody responses

To generate a broad and high-resolution view of the antibody response to human coronaviruses, including SARS-CoV-2, we designed and synthesized two separate DNA-barcoded 30mer peptide libraries, using the method described previously (Kozlov et al., 2012) (**Figure 1A**). Each library began as a pool of DNA oligonucleotide templates, which was modified using bulk enzymatic steps consisting of transcription, ligation of a puromycin-containing adapter oligo, translation, and reverse transcription. One library was focused on SARS-CoV-2 (‘SCV2’) and contained 2107 peptides representing the Spike and Nucleocapsid – the 2 most immunogenic coronavirus proteins – at high redundancy, with an average of 38 peptides covering each amino acid position (**Figure 1B**). The other library (human virome or ‘HV’) comprised 244,000 peptides designed to cover the full proteomes of all viruses known to infect humans, as of the end of 2018. Therefore, HV included peptides from the complete proteomes of 6/7 human coronaviruses: HCoV-229E, HCoV-OC43, HCoV-NL63, HCoV-HKU1, SARS-CoV, and MERS-CoV, but not SARS-CoV-2 (**Figure 1C**). The SCV2 library also included 373 positive control peptides that we have previously shown are commonly recognized across the human population (unpublished data). These controls represent a subset of the HV peptides and were designed from 55 different virus species.

**Figure 1.**
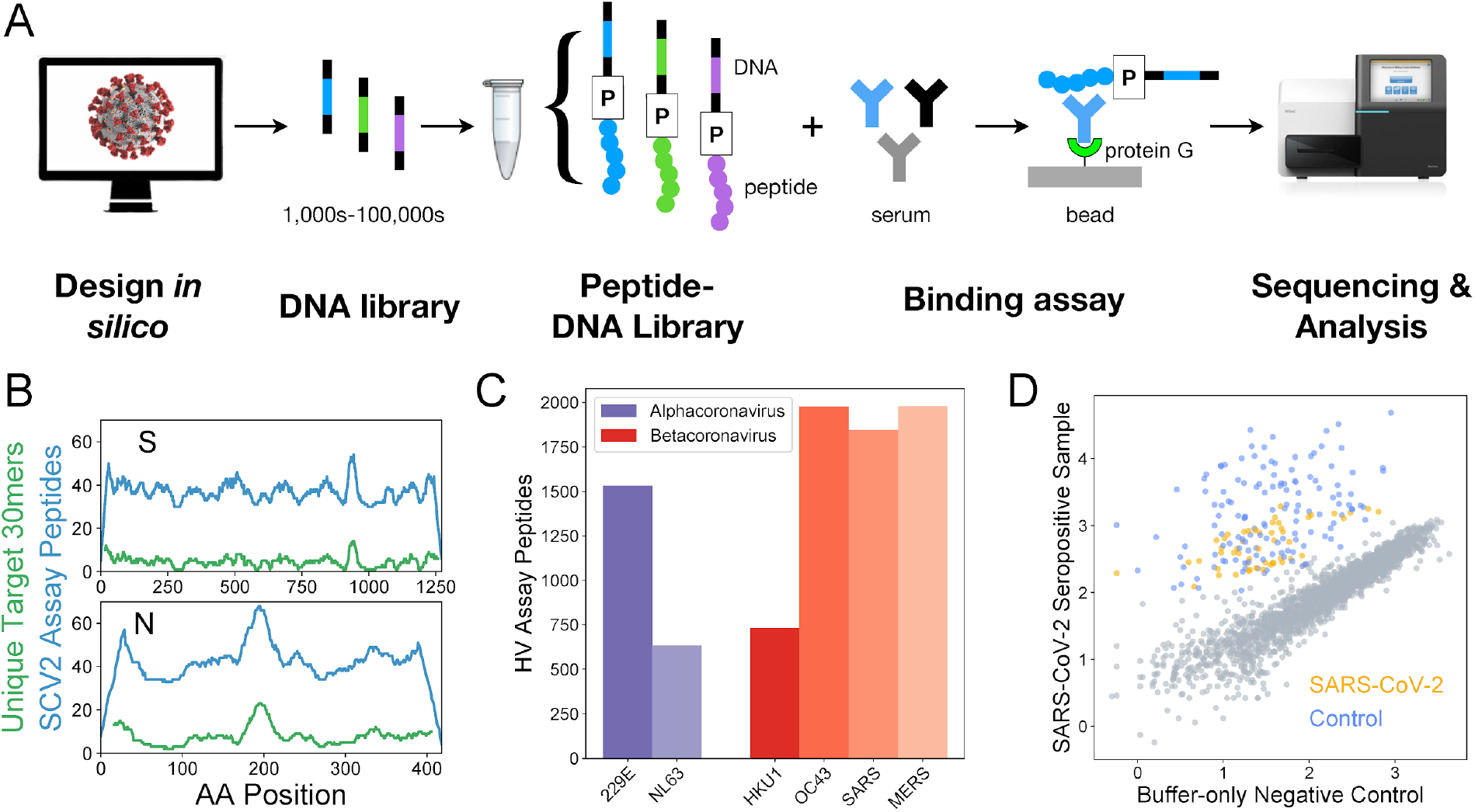
Epitope-resolved CoV serology using a highly-multiplexed peptide-based assay (‘PepSeq’). A) Platform for customizable highly-multiplexed peptide-based serology, comprising the following steps: (i) *in silico* design, (ii-iii) generation of a library of DNA-barcoded peptides from oligonucleotide templates using bulk *in vitro* reactions (transcription, ligation of a Puromycin (P)-containing adapter, translation, reverse transcription), (iv) serum binding assay and protein G capture, and (v) sequencing and analysis of the distribution of binders by their DNA barcodes. B) Peptide coverage depth across the SARS-CoV-2 spike (S) and nucleocapsid (N) proteins within the ‘SCV2’ peptide library. Peptide coverage depth (blue) correlates well with amino acid sequence diversity within the target SARS-CoV-2 sequences (green), calculated as the number of unique 30mers. C) Number of peptides within the HV library that were designed from each of the six human coronaviruses known prior to 2019. D) Example scatter plot illustrating SCV2 PepSeq assay results for a single serum sample. This plot shows normalized sequence read counts (log10 scale) for each peptide in the SCV2 library. Assay results using an antibody-free negative control are shown on the x-axis, while the results from a SARS-CoV-2 convalescent serum sample are shown on the y-axis. Grey circles represent unenriched peptides, with a strong correlation between the two assays, based on the starting abundance of the different peptides. Colored circles represent SARS-CoV-2 (orange) and non-SARS-CoV-2 control (blue) peptides that have been enriched through interaction with serum antibodies.

In total, we assayed and analyzed 27 COVID-19 convalescent and 21 SARS-CoV-2 negative (both pre- and post-pandemic) serum samples using our SCV2 PepSeq library (**Tables 1, S1**). Separately, we assayed 33 SARS-CoV-2 negative (pre-pandemic) serum samples using our HV PepSeq library. For each assay, we incubated our PepSeq probes overnight with serum/plasma (or buffer as a negative control), captured the IgG on protein G beads, washed away the non-binding library members, eluted binders, and then performed PCR and high-throughput sequencing on the DNA tags to identify the distribution of bound peptides. Each sample was run in duplicate, and we observed strong signal concordance between technical replicates of the same sera, including those run on different days (**Figure S1**). Comparative analysis of peptide abundance between serum/plasma and buffer-only negative controls revealed a strong correlation in abundance for the majority of peptides, while a subset of peptides showed distinctly higher relative abundance in each serum/plasma sample (**Figure 1D**). These latter peptides are those that have been enriched by binding to serum IgG. To quantify peptide enrichment, we calculated Z-scores for each peptide in each sample. For each peptide, relative abundance was normalized to the corresponding value for the buffer-only negative controls, and this normalized value was compared among peptides with similar abundance in the buffer-only controls. Each Z-score corresponds to the number of standard deviations away from the mean.

**Table 1.** Summary of samples characterized in this study.

| **PepSeq Library** | **Sample Type** | **Sample Size** | **Males/Females/Unreported** | **Median Age\*** | **Median days from diagnosis** |
| --- | --- | --- | --- | --- | --- |
| SCV2 | COVID-19 Convalescent | 27 | 14/13/0 | 51 | 26 |
|  | Negative Control | 21 | 11/6/4 | 37 | -- |
| HV | Negative Control | 33 | 0/0/33 | NA | -- |

NA = Not available, -- = Not applicable.
\*Median values were calculated from a subset of total samples for which this information was available. See Table S1 for details.

### Accurate detection of SARS-CoV-2 exposure and identification of epitopes

For the SCV2 PepSeq library, we evaluated the sensitivity/specificity for detection of SARS-CoV-2 exposure by generating receiver operating characteristic (ROC) curves with a sliding Z-score threshold and three different criteria for the number of enriched SARS-CoV-2 peptides needed for a positive result (**Figure 2B**). The SCV2 assay distinguished COVID-19 convalescent samples from negative controls with high accuracy (AUC = 0.89-0.92). Based on the ROC analysis and a qualitative assessment of the ability to discriminate signal from noise (**Figure 1C**), we selected a Z-score threshold of 11 for identifying enriched peptides; a peptide was required to meet or exceed this threshold in two technical replicates to be considered enriched. With only one SARS-CoV-2 peptide required for positivity, this threshold corresponded to a sensitivity of 81.5% and a specificity of 91.5%, with five false negative samples and two false positive samples. Notably, while both false positive samples exhibited at least one enriched peptide for both the N and S proteins, none of these peptides corresponded to the widely recognized, immunodominant epitopes observed for the COVID-19 convalescent sera (**Figure 2C**). To explore the potential for increasing sensitivity and specificity using a subset of SARS-CoV-2 peptides, we utilized a decision tree algorithm to identify the most discriminatory subset of peptides from our library. This analysis identified four SARS-CoV-2 peptides (indicated by green lines in **Figure 2C**) that were sufficient to detect all 22 convalescent donors that were called positive using the entire peptide set (**Table S2**). Using only these four peptides in the ROC analysis of all 48 donors increased the AUC to 0.97. With the same Z-score threshold of 11, the specificity increased to 100%, while sensitivity stayed at 81.5% (**Figure 2B**).

**Figure 2.**
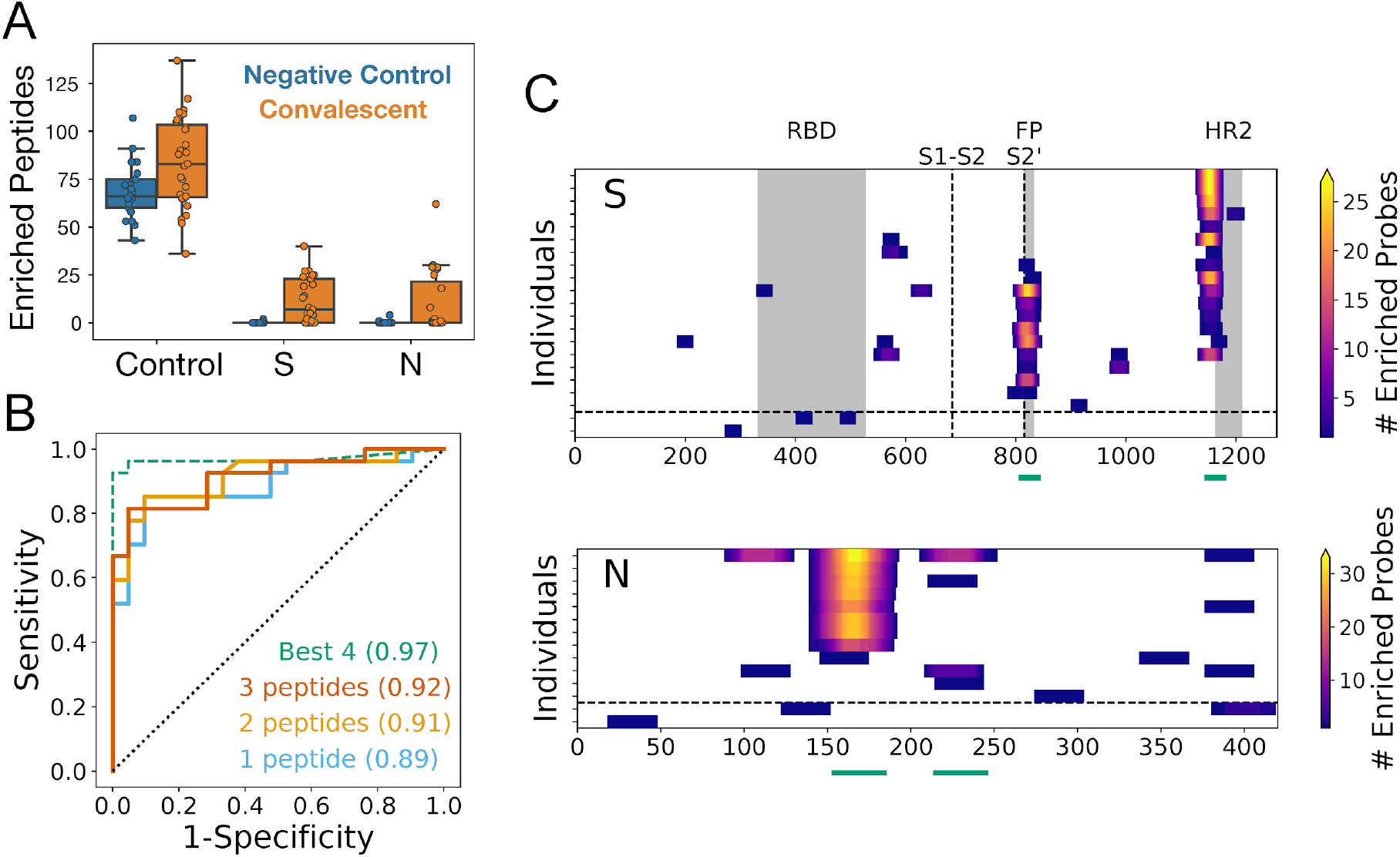
PepSeq identifies recurrent reactivities to SARS-CoV-2 peptides and classifies exposure status with high accuracy. A) Boxplots showing the number of enriched SCV2 library peptides from assays with negative control (blue, n=21) and SARS-CoV-2 convalescent (orange, n=27) samples, divided into three different categories: non-SARS-CoV-2 control peptides (Control), and SARS-CoV-2 Spike (S) and Nucleocapsid (N) peptides. All three of these comparisons are statistically significant (t-test, p<0.05). Individual data points are shown as circles, the limits of the boxes correspond to the 1st and 3rd quartiles, the black line inside each box corresponds to the median and the whiskers extend to points that lie within 1.5 interquartile ranges of the 1st and 3rd quartiles. B) ROC curves for prediction of SARS-CoV-2 exposure based on peptide-level Z-scores calculated for all SCV2 library peptides (solid lines) and for a subset of four peptides identified through a decision tree analysis (dashed line). Positivity of the assay was determined by the enrichment of peptides designed from SARS-CoV-2, and the full library analysis was run with three different thresholds for the number of enriched peptides required for a sample to be considered positive. For the analysis using only the “Best 4” peptides, only a single enriched peptide was required for a positive result. For all analyses, the AUC (shown in parentheses) was ≥0.89. C) Heat maps showing the locations of enriched SARS-CoV-2 peptides within the S and N proteins. Each row represents a single serum/plasma sample and each plot includes only samples with at least one enriched peptide. Each position is colored according to the number of enriched peptides that overlap that position. The horizontal dashed line separates COVID-19 convalescent samples (top) from negative control samples (bottom). The vertical dashed lines in the S protein plot represent the S1-S2 and S2’ cleavage sites, respectively. Grey boxes indicate selected functional regions: receptor binding domain (RBD), fusion peptide (FP) and heptad repeat 2 (HR2). Green lines below each plot indicate the positions of the “Best 4” peptides from panel (B).

As expected, multiple positive control peptides were found to be enriched in every serum sample that we tested (**Figure 2A**). Unexpectedly, we observed a small, but significant increase in the average number of enriched control peptides between convalescent and control donors, which involved peptides designed from a wide variety of virus species (t-test, p=0.01, 1.2 fold difference). However, this difference was small compared to the difference in the number of enriched SARS-CoV-2 peptides (56-fold, p=2e-5), and can likely be attributed to subtle differences in patient characteristics, sample collection, handling and/or storage among our different donor cohorts (****see Supplemental Results****). We do not expect this difference to impact our conclusions.

In total, we identified IgG reactivity (i.e., peptide enrichment) against 142 and 8 SARS-CoV-2 peptides in convalescent and negative control samples, respectively. There was no overlap between the reactive peptides observed in the convalescent and negative control samples (**Figure 2C**). The peptides enriched in convalescent samples clustered together into nine reactive regions of the S protein and six reactive regions of the N protein (**Figure 2C**), which together represent a minimum estimate for the number of epitopes (15). These epitopes were recognized at a range of prevalences across the sampled population. The most widely-recognized epitopes in S (positions 795-848 and 1127-1177) and N (positions 140-193) were each detected in 41-68% of the convalescent samples that tested positive with our assay (n=22) (**Figure 2C**), and >95% (21/22) of these convalescent samples were reactive to at least one of these three immunodominant regions. At the other extreme, 6 (43%) of the observed epitope regions were each detected in only a single donor. Despite the detection of a variety of SARS-CoV-2 S epitopes in the convalescent donors, very little reactivity was detected to peptides within the RBD, suggesting that these epitopes require protein conformations that are not well represented by linear 30mers.

To evaluate the potential for the identified S protein epitopes to be targeted by neutralizing antibodies, we evaluated these within the context of the protein’s structure. Of the S epitopes identified, four were recurrent across multiple convalescent samples, occurring at positions 1127-1177, 795-848, 543-589 and 971-1006 and in 14/27, 11/27, 4/27 and 2/27 convalescent donors, respectively. The enriched peptides at each of these four high-confidence regions were mapped onto a rendering of the recently-solved 3-dimensional structure of the native S trimer (**Figure 3A**). All four epitope regions are accessible for antibody binding on the surface of the trimer. The most widely-recognized reactive region (1127-1177) is located within the ‘stem helix’ just upstream and partially overlapping with the heptad repeat region 2 (HR2); this region is proximal to the transmembrane domain and partially unresolved in the native structure. The second epitope (795-848) resides at the S2’ cleavage site, spanning the fusion peptide whose exposure and incorporation into the host membrane are essential steps in virus entry into cells. The 543-589 and 971-1006 epitopes occur in the subdomain SD1 region (in the S1 subunit but C-terminal of the RBD), and the HR1 region of S2, respectively.. Comparison of pre- and post-fusion structures indicates that the HR2 epitope lies within a region that undergoes a dramatic conformational rearrangement during fusion (**Figure 3B,C**).

**Figure 3.**
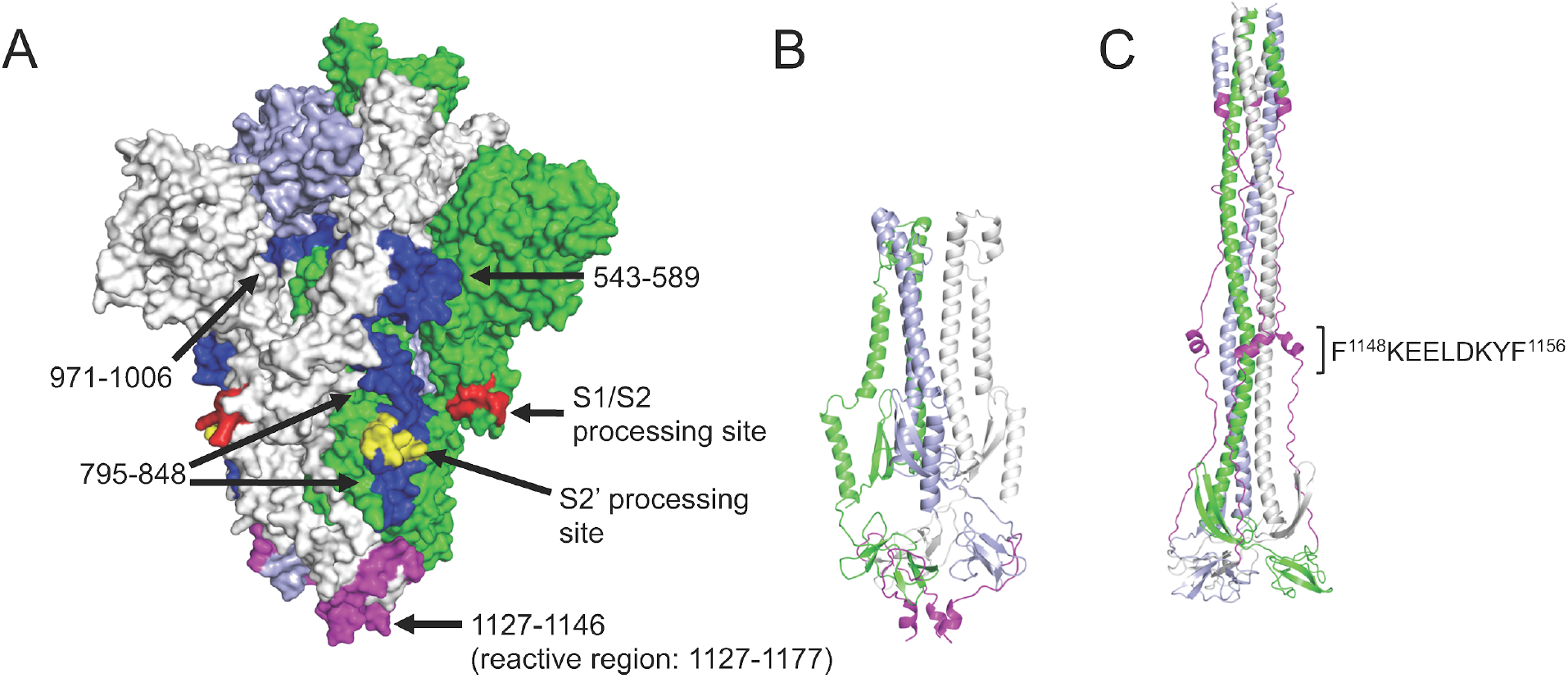
Recurrent Spike protein epitopes correspond to accessible and functionally-important sites within the protein structure. A) Space-filling SARS-CoV-2 Spike protein structure (PDB id: 6VYB) (Walls et al., 2020) showing the native trimer (monomers shown in green, gray and light blue) with the four recurrent epitope regions targeted by COVID-19 convalescent IgG (Figure 2C) highlighted in blue or magenta. Each epitope is identified by its amino acid range within the S protein sequence (GenBank: YP_009724390.1). The epitope at positions 1127-1177 (magenta) includes a region that is unresolved in the structure (1147-1177). Protease processing sites are highlighted in red and yellow, including the S2’ site that occurs within the 795-848 epitope. B) Ribbon model of the SARS-CoV-2 Spike S2 subunit after protease processing, using the same color scheme as in (A). C) Ribbon model of the 6-helical bundle (post-fusion) conformation of the S2 subunit, built based on the Cryo-EM structure of mouse hepatitis virus (PDB id: 6B3O) (Walls et al., 2017). The 1127-1177 region is again highlighted in magenta (with the minimal epitope region shown), and a comparison with (B) shows the dramatic conformational rearrangement that occurs at this site.

### Antibody epitopes and protein conservation across the human CoVs

To compare the SARS-CoV-2 reactivity profile described above with those of the other human coronaviruses, we performed a similar analysis but using the HV library (which covers all of the endemic human CoVs) and focusing on pre-pandemic donors. Applying the same Z-score threshold described above to the HV library, we identified reactivity to at least one endemic human coronavirus in 17 (51.5%) of the negative control samples we tested (n=33). To avoid false positives, we required ≥2 enriched peptides for a sample to be considered seropositive. Across all of the different coronaviruses, the vast majority of the recognized peptides were from the S and N proteins (95% of all enriched coronavirus peptides), with occasional reactivity observed to peptides in Orf1ab and a single peptide from the Membrane protein recognized in one sample (**Figure 4C**). Along with our SCV2 library data, these results indicate that S and N are the predominant antibody targets across all of the human-infecting coronaviruses and that pre-existing anti-CoV reactivity is common in the pre-pandemic population.

**Figure 4.**
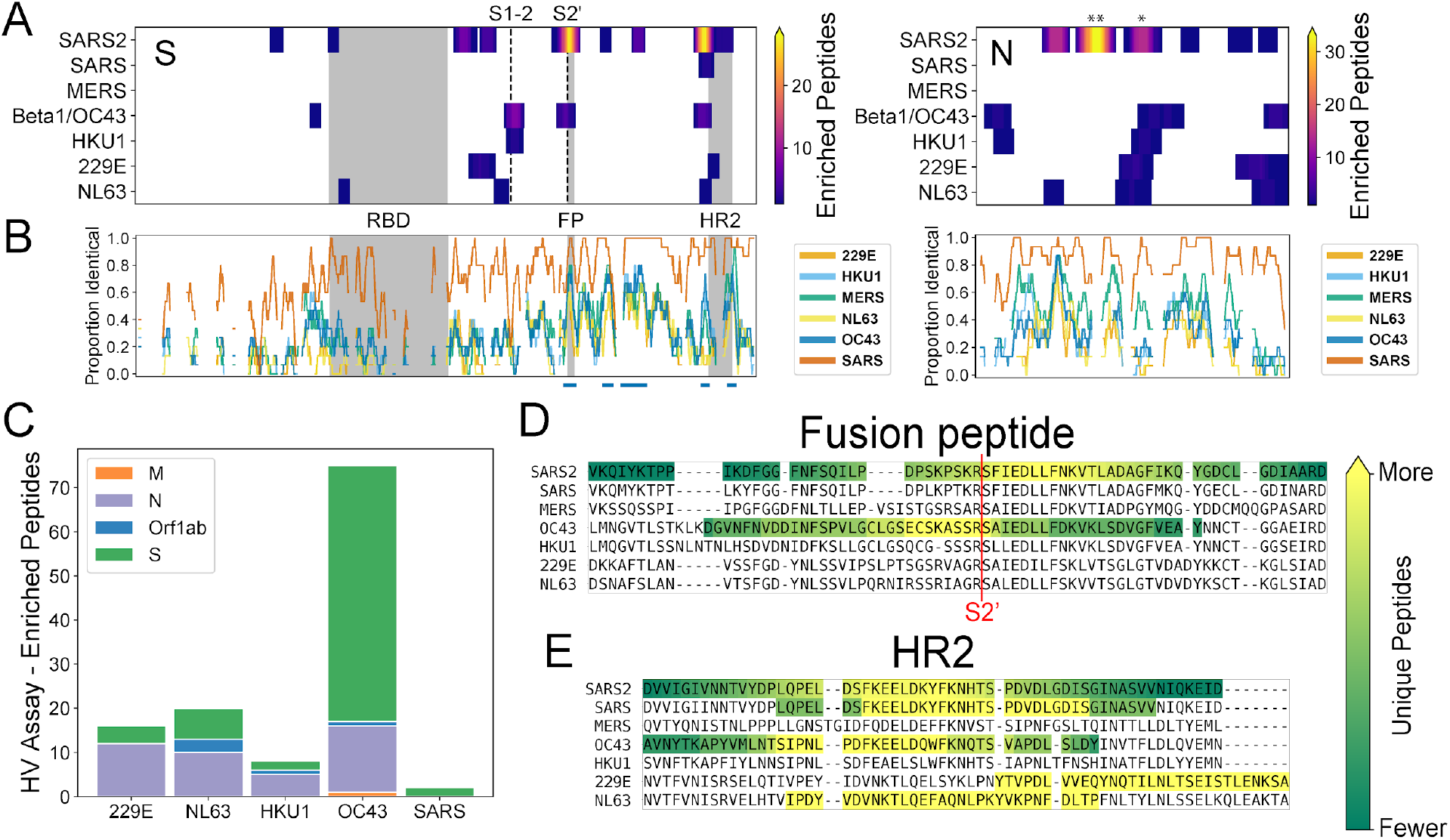
Recurrent SARS-CoV-2 epitopes correspond to conserved regions of Spike S2 that are also targeted in the response to other CoVs. A) Heat maps illustrating the relative locations of enriched SCV2 (from COVID-19 convalescent samples) and HV (from pre-pandemic controls) library peptides within the S (left) and N (right) proteins and across all human-infecting coronaviruses. Results have been aggregated across all tested samples and the color at each location indicates the number of unique enriched peptides. The vertical dashed lines in the S protein plot represent the S1-S2 and S2’ cleavage sites, respectively. Above the N plot, ‘**’ and ‘*’ indicate the 1st and 2nd most commonly immunogenic regions of this protein in COVID-19 convalescent samples, respectively. B) Comparison of amino acid sequence identity between SARS-CoV-2 and the other six human CoVs across the same S and N alignments used in panel (A). A sliding window of 15 amino acids was used and gaps represent windows with ≥30% indels. Blue bars under the S plot indicate regions ≥15 amino acids long that exhibit ≥70% identity between SARS-CoV-2 and hCoV-OC43 and/or hCoV-HKU1. Grey boxes in panels (A) and (B) indicate selected functional domains: receptor binding domain (RBD), fusion peptide (FP) and heptad repeat 2 (HR2). C) Protein-level distribution of enriched HV library peptides across five HCoVs and 33 pre-pandemic control samples. A single peptide could be counted multiple times if enrichment was independently observed in multiple samples. C-D) Multiple sequence alignments of the immunodominant and most widely-recognized protein regions of SARS-CoV-2, including representative sequences from each of the seven human coronaviruses. Regions containing enriched peptides are highlighted by colored backgrounds, with bright yellow indicating residues contained within the most unique enriched peptides and dark green indicating those contained within the least unique enriched peptides (colorbar scales are distinct among species). SARS-CoV-2 reactivity was determined using the SCV2 peptide library, while reactivity for the other coronaviruses was determined using the HV peptide library.

Within the S protein, we observed reactivity to homologous regions across multiple coronavirus species with highly variable percent identity to SARS-CoV-2 depending on the region and virus species (12.1 - 92.5% identical, average = 40%) (**Figure 4A**). Notably, we observed a correlation between amino acid sequence conservation among members of the Betacoronavirus genus and peptide enrichment in our assay. Across the full S protein, we identified five highly conserved regions (≥70% identical across 15mer sliding windows, blue bars in **Figure 4B**) between SARS-CoV-2 and each of the two endemic human betacoronaviruses: hCoV-OC43 and hCoV-HKU1 (four shared, one unique to each virus). All of these regions were located within the S2 subunit (**Figure 4B**), and while enriched SARS-CoV-2 peptides covered only 37% of the full S protein, we observed enriched peptides across almost all of the highly conserved regions: 96.4% (107/111) and 75.6% (93/123) of residues within these highly-conserved regions overlapped ≥1 enriched peptide for hCoV-OC43 and hCoV-HKU1, respectively.

Across the different coronavirus species, the most commonly recognized S protein region (‘HR2’) is also the most commonly reactive SARS-CoV-2 region within our convalescent donors. We detected reactivity in this region to 3/4 of the endemic human coronaviruses, though the precise locations of the recognized epitopes likely vary somewhat between species (**Figure 4E**). In one pre-pandemic serum sample, we also observed two enriched peptides in this region from the closely-related, epidemic-associated SARS-CoV species; however, these enrichments likely result from cross-reactivity with the endemic hCoV-OC43 (**Figure S2**, see below). For the Betacoronavirus 1 species (beta-CoV-1), which includes hCoV-OC43, we also detected reactivity at the same position as the second most immunodominant SARS-CoV-2 epitope, which overlaps the fusion peptide and S2’ cleavage site (**Figure 4A**). At this epitope region, we observed enrichment of one SARS-CoV peptide within a sample that also exhibited reactivity to homologous hCoV-OC43 peptides, again consistent with cross-reactive antibodies. However, the minimal epitope region contained within all enriched peptides is distinct between beta-CoV-1/hCoV-OC43 and SARS-CoV-2 (yellow residues in **Figure 4D**).

In contrast, we did not observe any reactivity for the endemic coronaviruses within the most commonly immunogenic SARS-CoV-2 N protein region (“**” in **Figure 4A**). However, we did observe homologous reactivities in other portions of the N protein. In fact the second most commonly immunogenic region observed in our COVID-19 convalescent samples (positions 206-252 in **Figure 2C**) overlaps with immunogenic regions in all four endemic human coronaviruses (“*” in **Figure 4A**). Somewhat surprisingly, however, we observed a somewhat greater similarity in the locations of reactive epitopes between SARS-CoV-2 and the endemic alphacoronaviruses (hCoV-229E and hCoV-NL63) than we did with the endemic betacoronaviruses (hCoV-HKU1 and hCoV-OC43) (**Figure 4A**).

### Interspecies cross-reactivity elicited by SARS-CoV-2 exposure

To explore the possibility that the antibody response to SARS-CoV-2 cross-reacts with other viruses, we focused on the panel of control peptides present in both the SCV2 and HV libraries (**Figure 5A**). This panel comprises 373 peptides from 55 virus species (range: 1-11 per species; 22 from the Coronaviridae family) for which we have previously observed recurrent reactivity in the general population. Consistent with previous results (not shown) and expected viral prevalences, we observed a range of positivity rates for these controls, including 25-100% for Rhinovirus-derived peptides and 0-48% for endemic human CoV-derived peptides. Comparing convalescent and negative control groups, Fisher’s exact tests identified a single peptide as significantly different between the groups at a Bonferroni-corrected threshold of p<1.3e-4 (**Figure 5A**). This peptide was enriched in 21/27 convalescents and 1/21 controls (p=2.5e-7), and was designed from a beta-CoV-1 strain. Although it was designed from a bovine coronavirus sequence, the peptide is 86.7% identical to the corresponding 30mer region in hCoV-OC43 (26/30 identical residues) and 100% identical to hCoV-OC43 across the 18 C-terminal residues. This peptide corresponds to positions 1218-1247 of the beta-CoV-1 S protein (UniProt: P36334) and it precisely overlaps the immunodominant HR2 region we identified based on SARS-CoV-2 peptides. Therefore, we hereafter refer to this peptide as ‘Beta1-HR2’. Beta1-HR2 also exhibits a high degree of conservation with SARS-CoV-2, particularly in the C-terminal portion of the peptide (66.7% identical across 18 C-terminal residues) (**Figure 5B**). By comparison, a second S-derived beta-CoV-1 peptide (‘Beta1-S12’) was reactive in about half of all samples tested (13/27 convalescents and 10/21 controls), indicating a high level of exposure to beta-CoV-1/hCoV-OC43 that did not differ between the groups (**Figure 5C**). Notably, the sequences of SARS-CoV-2 and beta-CoV-1 are highly divergent at the region covered by Beta1-S12 (13.3% identical) (**Figure 5B**).

**Figure 5.**
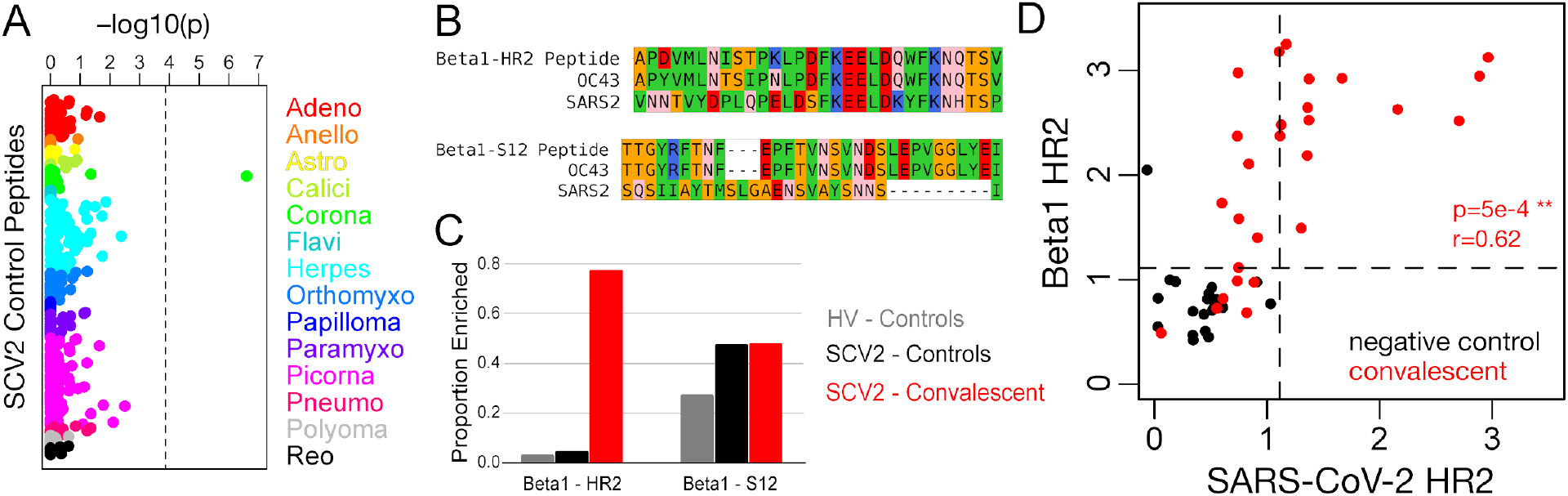
Spike HR2 antibodies elicited by SARS-CoV-2 strongly cross-react with the homologous region of Betacoronavirus 1. A) Fisher’s exact test p-values measuring the correlation between donor SARS-CoV-2 status and enrichment for each of 373 control peptides. These peptides were designed from 55 virus species that belong to 14 different families (colors, labels correspond to family names with the omission of “-viridae”), and they recognize epitopes that we previously identified as commonly reactive in the general population. Dashed vertical line shows the Bonferroni-corrected threshold for significance. B) Sequence alignments between SARS-CoV-2 (SARS2) and the Betacoronavirus 1 (beta1-CoV) strain, hCoV-OC43 (OC43), at two Spike protein regions covered by SCV2 library control peptides designed from beta1-CoV (Beta1) sequences. Residues are colored according to amino acid properties: small non-polar (orange), hydrophobic (green), polar (pink), negatively charged (red) and positively charged (blue). C) Proportion of samples reactive to the two Betacoronavirus 1 peptides shown in panel (B). Two separate sets of negative controls are shown, those assayed with the HV peptide library (grey, n=33) and those assayed with the SCV2 peptide library (black, n=21). Results from COVID-19 convalescent samples are shown in red (n=27). D) Quantitative comparison of reactivities to homologous HR2 peptides from SARS-CoV-2 (x-axis) and beta1-CoV (y-axis) across the SCV2-characterized donor cohort. Axes represent log10(2 + Z-scores) and dashed lines indicate threshold for significance (Z-score ≥ 11).

To further test the hypothesis that reactivity to Beta1-HR2 represents cross-reactivity with SARS-CoV-2, we compared donor-level reactivity to this peptide and a corresponding SARS-CoV-2 peptide (**Figure 5D**). We observed a significant positive correlation between measured reactivity against these two peptides in convalescent donors (r=0.62, p=5e-4), and all of the donors reactive to SARS-CoV-2-HR2 were also reactive to Beta1-HR2. However, an additional six convalescent donors were reactive to Beta1-CoV, despite a lack of reactivity to any SARS-CoV-2 peptides overlapping the HR2 epitope. Moreover, for donors reactive to either HR region, the signal strength for Beta1-HR2 was up to ~170-fold (mean ~10-fold) higher than for SARS-CoV-2-HR2, indicating that the anti-HR2 antibodies elicited by SARS-CoV-2 infection generally bind better to Beta1-HR2.

## Discussion

Like most viruses, SARS-CoV-2 elicits a robust antibody response whose targets are likely to be important determinants of disease outcome and the extent of protection conferred following natural infection or vaccination. In this study, we describe a customizable platform that enables epitope-resolved profiling of the antibody response (‘PepSeq’), and its application to the study of human CoVs including SARS-CoV-2. Using this system, we identify immunodominant epitopes in both the S and N proteins, several of which overlap conserved, functional sites in the Spike S2 subunit, and therefore have the potential to be sites of broadly neutralizing reactivity. By examining reactivity in pre-pandemic donors to homologous peptides from multiple human CoVs, we also show that the response to SARS-CoV-2 strongly cross-reacts with an endemic human CoV at one of these epitopes.

By independently testing reactivity across thousands of potential epitopes, we identified several with promise for use in both diagnostics and functional characterization assays. For two of the epitopes we detected in the S2 subunit of Spike – each discussed in further detail below – structural considerations, as well as previous characterization of related epitopes (Keng et al., 2005; Lai et al., 2005; Poh et al., 2020), strongly indicate neutralization potential. In these cases, a peptide-based assay may provide a facile means of profiling functional reactivities independently of cell/viral culture, and in a way that complements ACE2:RBD binding inhibition assays that cannot measure S2 reactivity (Tan et al., 2020). We also identified a set of 4 peptides across the S and N proteins that together exhibit great potential for generating an accurate profile of SARS-CoV-2 exposure. Although the precise diagnostic performance of this particular set needs to be quantified on a larger, independent sample set, our results provide a blueprint for a new generation of peptide-based diagnostics that would be easier to manufacture, and in some cases more informative, than existing full-protein/domain assays.

Our PepSeq analysis identified a novel epitope contained within positions 1127-1177 in Spike (minimal reactive sequence: FKEELDKYF) as the most widely-recognized SARS-CoV-2 linear epitope target in convalescent donors (**Figure 2C**). This region is located within the ‘stem helix’, directly N-terminal of the heptad-repeat 2 (HR2) region. While largely unresolved in the prefusion structure, analysis of post-fusion structures of CoV Spike proteins indicate that HR2 undergoes a ~180° reorientation during the formation of the 6-helix bundle in which it comes into close contact with the heptad-repeat 1 (HR1) region (Walls et al., 2017). HR-derived peptides that disrupt the HR1:HR2 interaction have previously been shown to inhibit infection by other CoVs (Liu et al., 2004; Xia et al., 2019), highlighting the strong potential for functional targeting of this region. Moreover, neutralizing monoclonal antibodies raised against related CoVs, including SARS-CoV (which has >95% amino acid-level identity to SARS-CoV-2 at the stem helix of HR2), have been shown to bind a region directly adjacent to the one that we identified in this study (Keng et al., 2005; Lai et al., 2005; Routledge et al., 1991). Strikingly, our analysis of reactivity across the human-infecting CoVs indicated that sites in the proximity of HR2 are also recognized in the responses to at least three of the four endemic species (**Figure 4A**). Since portions of this region are highly conserved across species (**Figure 4B**), cross-reactivity with pre-existing anti-CoV antibodies likely accounts for some of its immunodominance in the response to SARS-CoV-2, as discussed further below.

A second immunodominant reactivity that we identified in Spike S2 also occurs in a region whose sequence is highly-conserved across CoV species: positions 795-848 (minimal reactive sequence: EDLLFN), which overlaps the S2’ cleavage site and the Fusion Peptide (FP). Since the minimal region needed to explain the reactive peptides included residues on both sides of S2’ in many donors, this reactivity has the potential to block proteolytic processing and thereby prevent maturation of the S protein. Alternatively, and perhaps additionally, binding of antibody to the FP is expected to prevent its insertion into the host membrane and therefore prevent fusion and cell entry. A recent study, using a lower-throughput peptide-based approach also identified this FP epitope as reactive in two SARS-CoV-2 convalescent donors, and while they did not characterize the mechanism of action, they demonstrated the neutralization potential of antibodies against this epitope using antibody depletion assays (Poh et al., 2020). This study additionally reported an epitope downstream of the Spike RBD to which antibodies also exhibited neutralization potential. We observed reactivity to this same epitope in four of our COVID-19 convalescent donors (positions 543-589, minimal reactive sequence: LPFQQFGRDIADT). In addition to Spike S2 epitopes, we detected an immunodominant reactivity at positions 140-193 of the SARS-CoV-2 nucleocapsid protein, which lies at the C-terminal end of the domain that is primarily responsible for binding viral RNA (Chang et al., 2009). Unlike the reactivities described in Spike S2, this region does not appear to be targeted in the response to other CoVs (**Figure 4A**).

Despite well-documented serological reactivity in studies using the full-length RBD antigen (Amanat et al., 2020), we observed very little reactivity to peptides designed from the RBDs of human CoVs, including SARS-CoV-2 (Figures 2C, 3A). This lack of reactivity in our assay, as well as a similar absence of reactivity in a recent study using a lower-throughput peptide-based approach (Poh et al., 2020), suggests that antibodies to the RBD recognize conformational epitopes and/or depend on post-translational modifications. Like other peptide-based antibody assays, PepSeq is limited to the detection of epitopes that are well-represented by linear peptides and do not require post-translational modifications. The dependence of RBD epitopes on secondary/tertiary structure is supported by structural analyses of the footprints of neutralizing antibodies bound to Spike RBD, which indicate the involvement of residues that are distal in the linear sequence (Pinto et al., 2020; Yuan et al., 2020). The identification of epitopes like these will require lower throughput approaches including mutagenesis and/or structural studies.

The observation that ~80% of SARS-CoV-2 convalescent donors react strongly to a Beta1-HR2 peptide targeted in ~5% of our negative control samples (**Figure 5C**) is, to our knowledge, the first identification of a B cell epitope for which there is cross-reactivity between the pandemic virus and an endemic pathogen. The fact that antibodies against Beta1-HR2 occur in individuals who also have antibodies targeting SARS-CoV-2-HR2, but with, on average, ~10X greater signal strength, is most consistent with a model in which pre-existing B cell clones raised against hCoV-OC43 are recruited into the response to SARS-CoV-2. In further support of this hypothesis, the one pre-pandemic donor in which we observed a strong Beta1-HR2 response with our HV assay also exhibited reactivity to two HR2 peptides designed from SARS-CoV (no SARS-CoV-2 peptides are present in our HV library) (**Figure S2**). Pre-existing cross-reactive clones would be expected to have a range of intrinsic affinities for the homologous SARS-CoV-2 epitope, and these could be further improved by somatic mutation. However, by analogy with other viruses, the fact that presumed exposure to hCoV-OC43 precedes exposure to SARS-CoV-2 may limit the efficiency with which the response can be redirected, due to ‘imprinting’ (Gostic et al., 2016; Monto et al., 2017), which could account for the systematic difference in affinities to the corresponding epitopes from the two species. Under this model, the ~20% of convalescent donors who exhibit detectable reactivity to Beta1-HR2 but not to SARS-CoV-2-HR2 (upper left quadrant of Figure 5D) represent cases where pre-existing antibodies to hCoV-OC43 bind only weakly to SARS-CoV-2 (below the threshold of the PepSeq assay) and have been unable to acquire a high affinity against the new virus. This model also suggests that anti-Beta1-HR2 B cell memory that is capable of cross-reacting with SARS-CoV-2 is prevalent in the general population – consistent with the near universal seropositivity reported for HCoV-OC43 (Gorse et al., 2010) – although often below our limit of detection.

Our findings raise the possibility that the nature of an individual’s antibody response to prior hCoV-OC43 infection may impact the course of COVID-19 disease. They also indicate that analysis of S2 reactivity is crucial for a complete assessment of the humoral response to SARS-CoV-2, consistent with the observation that S2-only assays provide an equally strong correlate of neutralization compared to RBD-only assays (J. Nikolich and D. Bhattacharya, personal communication). The HR2 cross-reactivity characterized here represents a possible source of background reactivity for SARS-CoV-2 serological assays that include the S2 subunit of Spike, which would be absent in those targeting only the RBD, for which sequence conservation is lower across CoV species (Khan et al., 2020). However, our findings also indicate that the incorporation of related beta-CoV antigens may improve the sensitivity of SARS-CoV-2 serological analyses, and in particular, that a differential analysis of SARS-CoV-2 and hCoV-OC43 Spike reactivity may provide an important measure of the efficiency with which pre-existing cross-reactive responses can be redirected. Furthermore, based on the level of sequence conservation at the S2’ cleavage/fusion peptide site, we expect that similar cross-reactivity may also occur at this site, and, in fact, we observed preliminary evidence for such cross-reactivity in one of the pre-pandemic controls analyzed with our HV library (**Figure S2**). However, due to the absence of an hCoV-OC43 S2’ control peptide in our SCV2 library, we were not able to directly evaluate the potential for this cross-reactivity in COVID-19 convalescent donors.

The identification of broadly-immunogenic epitopes in conserved functional domains of the SARS-CoV-2 Spike S2 subunit, including cross-reactivity with an endemic human CoV, also has implications for the design of therapeutic antibodies and vaccines. SARS-CoV-2 vaccines currently under development predominantly use two forms of the S antigen – whole protein or the RBD – and in each case are designed primarily to elicit neutralizing antibodies (WHO, 2020). Relative to RBD-focused vaccines, we hypothesize that vaccines that include the Spike HR2 and FP sites: (i) will be able to induce a broader array of neutralizing reactivities, (ii) may be more capable of rapidly recruiting pre-existing memory B cells that are prevalent in the population and (iii) may be less prone to viral escape due to a lower tolerance for amino acid substitutions. In particular, the identification of HR2 as a conserved, functionally-important and broadly-immunogenic site capable of eliciting cross-reacting antibodies, makes this region a candidate for the development of broadly-neutralizing responses against betacoronaviruses.

## Methods

### Samples

COVID-19 convalescent serum and plasma samples (n=27) were collected at sites in California, USA (Vitalant Research Institute and City of Hope National Medical Center) and Norway (St Olav’s Hospital, Trondheim and Oslo University Hospital, Oslo) from patients who had tested positive for SARS-CoV-2 by RT-PCR a median of 26 days prior (see **Table S1** for details). Post-pandemic negative control samples (n=4) were also collected from healthy donors at one of these sites (City of Hope National Medical Center). All post-pandemic sample collections and use proceeded in compliance with the relevant regulatory requirements, including IRB approval where appropriate. Pre-pandemic negative control serum samples characterized using our SCV2 library (n=17) were provided by Creative Testing Solutions (Phoenix, AZ). These samples were collected during January 2015 from multiple locations in California (**Table S1**). Pre-pandemic negative control serum samples characterized using our HV library (n=33) were collected at Walter Reed National Military Medical Center (Bethesda, MD) during 2019 (latest collections were during the first week of December). The use of all pre-pandemic samples was reviewed by the NAU and TGen IRB Offices and determined not to be human subjects research.

### PepSeq Library Design

We designed two different libraries of peptides in order to assess antibody reactivity to SARS-CoV-2 peptides and to peptides from other human-infecting coronaviruses. The first set of peptides (Human Virome, HV) was designed to broadly cover potential epitope diversity for all viruses known to infect humans. To generate this design, we downloaded all protein sequences available in UniProt, on November 19, 2018, that were linked to 474 viral species-level taxonomy IDs (see Supplemental Materials for details). Following a series of quality filters to remove identical sequences, those that were too short (<30 aa), those that contained recombinant non-viral sequences and those that were taxonomically misclassified (see Supplemental Materials), we were left with 1,300,994 target protein sequences. In order to control for sampling bias within the database, we randomly subsampled overrepresented virus species, including no more than 2000 and 4000 sequences for viruses with RNA and DNA genomes, respectively. Additional protein sequences were allowed for DNA viruses because they often contain larger genomes and proteomes (i.e., more distinct genes). When down-sampling, priority was given to proteins from the Swiss-Prot database, which have been manually reviewed. Our final down-sampled target set included 148,215 protein sequences and 88.78 M amino acids.

Our HV peptides were designed using an epitope-centric set cover design algorithm (https://github.com/LadnerLab/Library-Design), with a focus on optimizing 9mer (i.e., 9 amino acids long) epitope coverage using 30mer peptides. To reduce the runtime and memory requirements of the algorithm, we partitioned our target protein sequences according to taxonomy. Given the high levels of genetic divergence between viral families and genera, we do not expect that this partitioning substantially impacted our final design. Including a small set of negative control peptides selected from eukaryotic proteins, this design included 244,000 unique 30mer peptides, and represents approximately 70% of all potential 9mer epitopes contained within the target protein sequences. Each of these peptides was represented by a single nucleotide encoding. This design does not contain any peptides derived from SARS-CoV-2, but does contain full proteome coverage of the other six coronaviruses known to infect humans: Human coronavirus 229E (NCBI taxID: 11137), Human coronavirus NL63 (277944), Human coronavirus HKU1 (290028), Betacoronavirus 1 (694003, includes Human coronavirus OC43), Severe acute respiratory syndrome-related coronavirus (694009, SARS), and Middle East respiratory syndrome-related coronavirus (1335626, MERS).

Our second design (SCV2) focused almost entirely on SARS-CoV-2, including high density tiling of peptides across the two most immunogenic SARS-CoV-2 proteins: the spike glycoprotein (S) and the nucleocapsid protein (N). As targets for this design, we utilized 2303 SARS-CoV-2 genome sequences downloaded from GISAID on April 3rd, 2020 (Table S3), along with six locally generated sequences. Using these genomes, we first generated consensus amino acid sequences for the S and N proteins. In our design, we included all of the unique 30mer peptides contained in these consensus sequences, equivalent to a 1-step sliding window approach (Shiryaev et al., 2012). Additionally, we used the same epitope-centric set cover design algorithm used for HV in order to capture amino acid-level polymorphisms present within our full set of target genomes. This aspect of the design ensured that 100% of the unique 16mer peptides present in the S and N proteins from the 2309 SARS-CoV-2 genomes were represented in our design. In total, this design included 1550 30mer peptides from the S protein and 557 30mer peptides from the N protein. Each of these peptides was represented by three different nucleotide encodings. This design also included a set of 373 control peptides. These controls represent a subset of the HV peptides, which we have determined are commonly recognized by IgG antibodies in human sera (unpublished results). Therefore, we expect that some fraction of these controls will be recognized by antibodies in each blood sample tested. Collectively, these peptides were designed from 55 different virus species, including the four endemic human coronaviruses.

### PepSeq Library Synthesis and Assay

Libraries of covalently-coupled peptide:DNA conjugates were prepared from pools of DNA oligonucleotide templates in bulk enzymatic reactions using the method described previously (Kozlov et al., 2012), with minor modifications as noted hereafter. Briefly, pools of ssDNA templates (*Agilent*) were PCR-amplified and the dsDNA products were used as templates for *in vitro* transcription (*Ampliscribe*). The resulting mRNA was ligated to a hairpin oligonucleotide adapter bearing a puromycin molecule tethered by a PEG spacer and, following buffer exchange, the reaction mix was used as a template in an *in vitro* translation reaction (*PURExpress, NEB*). Constructs bearing mRNA – comprising of (i) mRNA, (ii) mRNA+adapter, (iii) mRNA+adapter+peptide – were isolated using magnetic beads coated with a DNA oligo complementary to a 30-mer sequence in the mRNA constant region. A reverse transcription reaction, primed by the adapter hairpin, was used to generate cDNA, after which RNase was applied to remove mRNA. Product was buffer-exchanged, quantified by running on a gel against standard DNA oligos of known concentrations, and used without further modifications or purification.

To perform serological assays, 5uL of a 1:10 dilution of serum/plasma in Superblock T20 (*Thermo*) was added to 0.1pmol of PepSeq library for a total volume of 10uL and was incubated at 20℃ overnight. The binding reaction was applied to pre-washed protein G-bearing beads (*Thermo*) for 15 minutes, after which beads were washed 10 times with 1x PBST. After the final wash, beads were resuspended in 30uL of water and heated to 95℃ for 5 minutes to elute bound product. Elutions were amplified and indexed using barcoded DNA oligos. Following PCR cleanup, products were pooled, quantified and sequenced on a NextSeq instrument (*Illumina*).

### PepSeq Data Analysis

We used PepSIRF v1.3.0 (Fink et al., 2020), along with custom scripts (https://github.com/LadnerLab/PepSIRF/tree/master/extensions), to analyze the PepSeq HTS data. The data analysis included three primary steps: 1) demultiplexing and assignment of reads to peptides, 2) calculation of enrichment Z-scores individually for each assay and peptide and 3) identification of enriched peptides for each sample based on the consistency of Z-scores across replicates.

Demultiplexing and assignment of reads to peptides was done using the *demux* module of PepSIRF (Fink et al., 2020), allowing up to 1 mismatch within each of the index sequences and up to 2 mismatches with the expected DNA tag (90 nt in length). Z-scores were calculated using a method adapted from (Mina et al., 2019). This process involved the generation of peptide bins, each of which contained ≥300 peptides with similar starting abundance in our PepSeq assay. Starting abundance for each peptide was estimated using buffer-only controls. In total, 4-8 independent buffer-only controls were used to generate the bins for this study. The raw read counts from each of these controls were first normalized to reads per million (RPM) using the column sum normalization method in the *norm* module of PepSIRF. This was to ensure that independent assays were weighted evenly, regardless of differences in the depth of sequencing. Bins were then generated using the *bin* PepSIRF module.

Z-scores were calculated using the *zscore.py* script, and each Z-score corresponds to the number of standard deviations away from the mean, with the mean and standard deviation calculated independently for the peptides from each bin. It is important that the mean and standard deviation reflect the distribution of unenriched peptides within a bin. Therefore, these calculations were based on the 75% highest density interval of read counts within each bin. Prior to Z-score calculation, RPM counts for each peptide were further normalized by subtracting the average RPM count observed within our superblock-only controls. This second normalization step controlled for variability in peptide starting abundance within a bin. Finally, the *p_enrich* module of PepSIRF was used to determine which peptides had been enriched through our assay. This module identifies peptides that meet or exceed minimum thresholds, in both replicates, for Z-score and RPM count. Decision tree analysis was conducted using the DecisionTreeClassifier() method in the Scikit-learn Python module, v0.20.1.

### Visualization of protein structure

To visualize our identified SARS-CoV-2 epitopes within the 3D conformational structure of the S protein, we utilized the cryo-electron microscopy (Cryo-EM) structure available in the RCSB Protein Data Bank (PDB id: 6VY). To compare epitope positions across CoV species, we built three additional structures using Cryo-EM templates from PDB: 5SZS for hCoV-NL63, 6ACD for SARS-CoV and 6NZK for hCoV-OC43. We performed structural modelling using Swiss-Model software (Waterhouse et al., 2018). Structural alignments and image preparation were done with PyMOL (version 2.3.2, Schrodinger, LLC). To build models of the post-fusion state for S2 subunit fragments, we used the Cryo-EM structure for mouse hepatitis virus, determined by Walls et al. (PDB id: 6B3O) (Walls et al., 2017).

## Acknowledgements

This work was supported by the National Institute of Allergy and Infectious Diseases of the National Institutes of Health under Award Numbers U24AI152172 and U24AI152172-01S1, the National Institute on Minority Health and Health Disparities of the National Institutes of Health under Award Number U54MD012388 and the State of Arizona Technology and Research Initiative Fund (TRIF), administered by the Arizona Board of Regents, through Northern Arizona University. The content is solely the responsibility of the authors and does not necessarily represent the official views of the National Institutes of Health. ZWF was also supported by a Hooper Undergraduate Research Award and a Jean Shuler Research Mini-Grant. We are grateful for assistance from Mike Busch and Phillip Williamson in obtaining samples and Piotr Swiderski in synthesizing a custom oligonucleotide reagent. We are also grateful for the constructive feedback received from Paul Keim, Stephen Daley, Gerard Zurawski and Erik Settles, which greatly improved the manuscript.

## Author Contributions

Categories are as defined by CRediT taxonomy. Authors are listed in the same order as the byline.

Conceptualization: JTL, JAA; Methodology: JTL, NM, MSC, JAA; Software: JTL, ZWF; Formal Analysis: JTL, JAA; Investigation: SNH, ASB, ALE, FR; Resources: JD, KES, MS, WD, SD, JY, MAC, MB, MHF, SAN, DEK; Data Curation: JTL, ZWK, SAS; Writing – Original Draft: JTL, JAA; Writing – Review & Editing: SNH, ASB, ALE, ZWF, FR, DEK, SAS; Visualization: JTL, SAS, JAA; Supervision: JTL, JAA; Project Administration: JTL, JAA; Funding Acquisition: JTL, JAA.

## Notes

http://github.com/jtladner/Manuscripts/tree/master/2020_Ladner_SARS2PepSeq

### Competing Interest Statement

The authors have declared no competing interest.

