## Supplemental Material for "Epitope-resolved profiling of the SARS-CoV-2 antibody response identifies cross-reactivity with an endemic human CoV"

### Supplemental Results

#### Enriched Control Peptides

We observed a small, but significant increase in the average number of enriched SCV2 library control peptides between COVID-19 convalescent and negative control donors (t-test, p=0.01, 1.2 fold difference) (**Figure 2A**). However, this difference was small compared to the difference in the number of enriched SARS-CoV-2 peptides (56-fold, p=2e-5). There was a significant difference in age between our convalescent and negative control donors, with our negative control donors being slightly younger on average than our COVID-19 convalescent donors (**Figure S3A**, 1.3-fold, p=0.015). However, within our sample set, we did not observe a correlation between donor age and the number of enriched control peptides (**Figure S3B)**. In contrast, we did observe several significant pairwise differences in the number of enriched control peptides when we compared samples obtained from different sources (**Figure S3C**). Specifically, we observed significantly fewer enriched control peptides in our pre-pandemic negative controls (1.2-1.5-fold, p=0.001-0.01). Therefore, this difference can likely be attributed to subtle differences in patient characteristics, sample collection, handling and/or storage among our different donor cohorts. However, we cannot rule out a role of recent SARS-CoV-2 infection in boosting overall antibody responses in convalescent donors.

### Supplemental Methods

#### HV Library Design

For our human virome ('HV') peptide library, we sought to include sequences from all viruses known to infect humans. For viruses with RNA genomes, we obtained a list of 214 virus species from (Woolhouse and Brierley, 2018). NCBI taxonomy IDs were obtained for each of these species using the "names.dmp" file from the NCBI "new_taxdump" downloaded on 11-19-2018 [note: "Bovine viral diarrhea virus 1" (NCBI:txid11099) was replaced with the corresponding species, "Pestivirus A" (NCBI:txid2170080)]. Taxonomy IDs for human viruses with DNA genomes were obtained using the "host.dmp", "nodes.dmp" and "fullnamelineage.dmp" files from the NCBI "new_taxdump" downloaded on 11-26-2018. In total, we identified 289 taxonomy IDs annotated as virus species with DNA genomes that are known to cause human infections; however, 31 of these were excluded from our design because they clearly belonged to unclassified adenovirus strains, rather than distinct virus species. Finally, we included two taxonomy IDs associated with the Jingmenvirus group, members of which have recently been associated with human infections in China (Jia et al., 2019).

On November 19, 2018, we downloaded all viral protein sequences from the UniProt Knowledgebase ("uniprot_sprot_viruses.dat" and "uniprot_trembl_viruses.dat" from ftp://ftp.uniprot.org/pub/databases/uniprot/current_release/knowledgebase/taxonomic_divisions/, and extracted the sequences annotated with one of our 474 target species taxonomy IDs. NCBI BLAST was used to identify sequences with non-viral components (i.e. recombinant), specifically those containing common reporter and therapeutic proteins: ubiquitin, luciferase, green fluorescent protein, chloramphenicol acetyltransferase, LacZ, GusA and GusB. These sequences were excluded from the assay design. To identify taxonomically misclassified proteins, we downloaded all of the proteins annotated in the NCBI RefSeq database for our target species, when available (342/474 target species IDs). We then used NCBI BLAST to identify the best matching RefSeq protein for each UniProt protein, and flagged instances when the top hit was "strong" and to a RefSeq protein from a different genus (≥80% nt identity) or species (≥95% nt identity). All of the flagged UniProt proteins were manually investigated, including an additional BLAST to the NCBI nt database, and sequences confirmed to be misclassified were either removed completely or taxonomically relabeled. Finally, we removed all sequences <30 amino acids in length and collapsed identical sequences to a single representative using a custom python script (https://github.com/LadnerLab/Library-Design/blob/master/scripts/one_hundred_rep.py).

Following our length, identity and taxonomy filters, we were left with 1,300,994 target protein sequences assigned to 443 distinct species taxonomy IDs. However, a small subset of viral species contributed the vast majority of protein sequences. For example, 49% of the proteins were from human immunodeficiency virus 1 and 16% were from influenza A virus. To ensure more even representation of viruses within our assay design, we randomly subsampled the overrepresented species, including no more than 2000 and 4000 proteins for viruses with RNA and DNA genomes, respectively. Additional protein sequences were allowed for DNA viruses because they often contain larger genomes and proteomes (i.e., more distinct genes). When down-sampling, priority was given to proteins from the Swiss-Prot database, which have been manually reviewed. Our final down-sampled target set included 148,215 proteins and 88.78 M amino acids.

In order to optimize potential epitope coverage in as few peptides as possible, we utilized a greedy set cover algorithm in which all potential linear epitopes contained within our target sequences were treated as our "elements of interest" and "sets" were defined as the collection of all potential epitopes contained within a potential peptide probe. Each round, a score was calculated for each potential peptide probe, which corresponded to the sum of the frequencies of each contained epitope within the full target set of proteins, and the highest scoring peptide was added to our design. In the event of a tie, a peptide was randomly chosen from the highest scoring subset. All of the epitopes contained within the added peptide were then excluded from the calculation of scores in the next round. This procedure was repeated until a targeted proportion of total epitope diversity was contained within the selected peptides. This algorithm was implemented with custom software (https://github.com/LadnerLab/Library-Design). For our design, we focused on optimizing 9mer (i.e., 9 amino acids long) epitope coverage using 30mer peptides.

To reduce the runtime and memory requirements of the algorithm, we partitioned our target protein sequences according to taxonomy prior to running our peptide design algorithm. We generated subsets of our target proteins by first dividing according to viral family and finally by genus, if the family-level partition contained >500,000 unique 9mers. Due to the random nature of peptide selection in the event of a tie, our algorithm is not deterministic. Therefore, we independently ran the design for each partition 5-20 times (depending on the size of the partition), and in each case, we selected the result with the fewest number of chosen peptides.

For a subset of species with low numbers of UniProt sequences per annotated protein, we added unique protein sequences present in GenBank to our list of targets. Additionally, for these species and one other with low overall epitope coverage in our set cover design (severe fever with thrombocytopenia syndrome virus, taxID=1933190), we redesigned peptides using a sequence-level (i.e., no alignment) sliding window approach (step=19) in order to optimize epitope coverage. We also included 15 “positive control” peptides, which included epitopes known to be broadly reactive in the human population based on preliminary, unpublished data, and 223 “negative control” peptides designed from an assortment of eukaryotic proteins of exotic species (e.g., coelacanth, coral, great white shark).

### Supplemental Figures

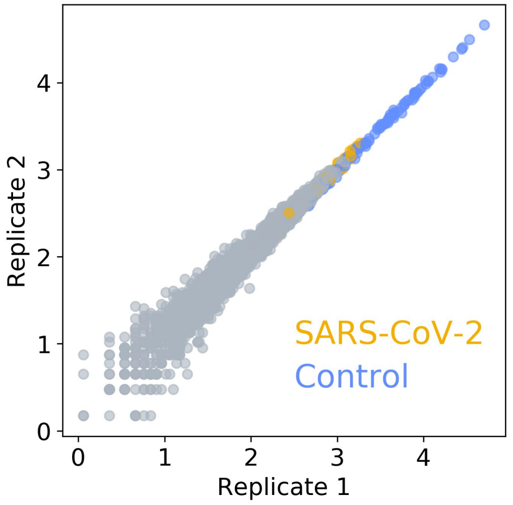


**Figure S1. Strong correlation between replicate PepSeq assays run separately on the same serum sample.** Axes show normalized read counts (log10 scale) for each peptide in the SCV2 library. Grey circles represent unenriched peptides. Colored circles represent SARS-CoV-2 (orange) and non-SARS-CoV-2 control (blue) peptides that have been enriched through interaction with serum antibodies.

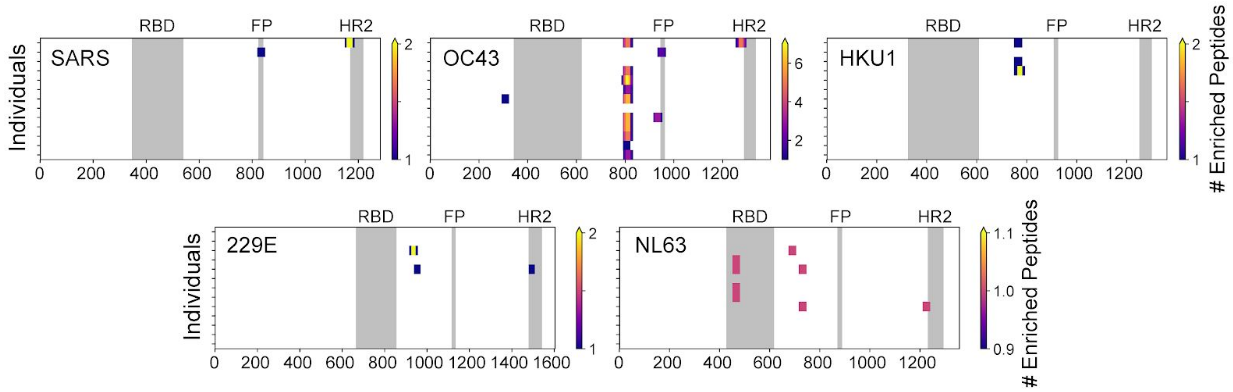


**Figure S2. Distribution of enriched peptides from CoV Spike proteins in the HV library.** Each row represents a pre-pandemic negative control sample that was determined to be seropositive for at least one of the non-SARS-CoV-2 human infecting coronaviruses (i.e., enrichment of ≥2 peptides from a non-SARS-CoV-2 coronavirus). The same 13 samples are shown in the same order in each plot. The focal coronavirus species is indicated in the top left corner of each plot: SARS-CoV ('SARS'), Beta1-CoV/hCoV-OC43 ('OC43'), hCoV-HKU1 ('HKU1'), hCoV-229E ('229E'), and hCoV-NL63 ('NL63'). Each position is colored according to the number of enriched peptides that overlap that position. Grey boxes indicate selected functional regions: receptor binding domain (RBD), fusion peptide (FP) and heptad repeat 2 (HR2). Both samples exhibiting reactivity to SARS-CoV peptides (top two rows) also exhibit hCoV-OC43 reactivity in homologous regions, consistent with cross-reactivity between peptides derived from endemic and epidemic coronavirus species. Both serum samples exhibiting reactivity to SARS-CoV peptides were collected in 2019 (16 years after the SARS-CoV epidemic) in Bethesda, MD, USA. Given the timing of these samples and the very small number of documented SARS-CoV cases in the US (Centers for Disease Control and Prevention (CDC), 2003), it is highly unlikely that these individuals have actually been exposed to SARS-CoV.

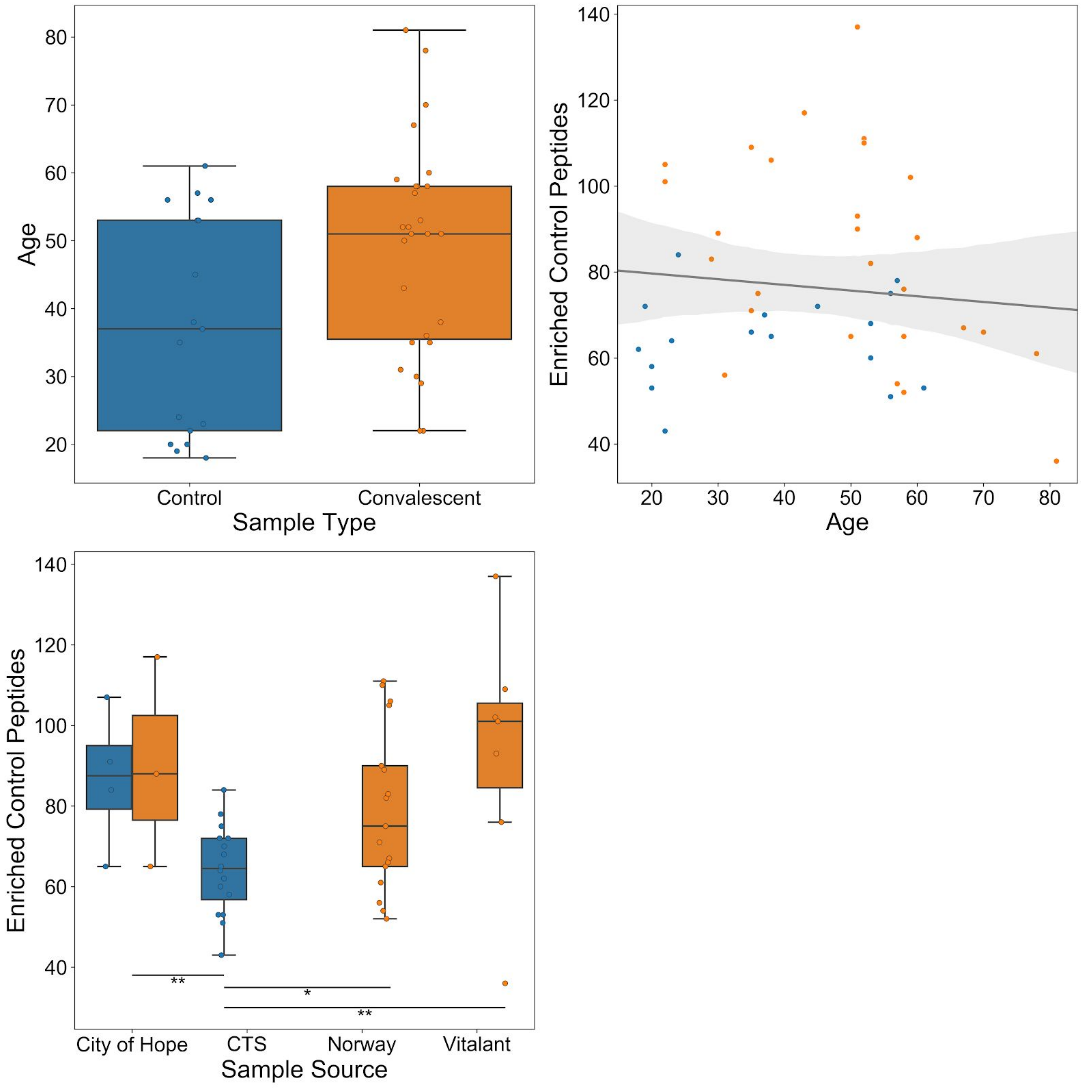


**Figure S3. Effect of age and sample source on number of enriched control peptides.** A) Boxplots depicting donor age distributions for negative control and convalescent serum/plasma samples. The means of these distributions are significantly different based on a t-test (p=0.022). B) Scatterplot comparing donor age (x-axis) and the number of enriched SCV2 library control peptides (y-axis). Each circle represents a single serum/plasma sample. Grey line and band represent the best fit linear regression line and 95% confidence interval, respectively, as estimated by the Seaborn regplot() function. C) Boxplots depicting SCV2 library enriched control peptide distributions for each sample source and type. The mean for the negative control samples from Creative Testing Solutions (CTS) is significantly lower than that for the samples from the three other sources based on t-tests. Significantly different pairs are indicated with horizontal lines: *<0.05, **<0.01. For all boxplots, individual data points are shown as circles, the limits of the colored boxes correspond to the 1st and 3rd quartiles, the black line inside each box corresponds to the median and the whiskers extend to points that lie within 1.5 interquartile ranges of the 1st and 3rd quartiles. In all panels, blue is used to indicate negative control samples and orange for convalescent samples.

### Supplemental Tables

**Table S1. Metadata for clinical samples assayed with SCV2 library peptides.**

Available at github.com/jtladner/Manuscripts/tree/master/2020_Ladner_SARS2PepSeq.

**Table S2. SARS-CoV-2 peptides chosen by decision tree algorithm for discriminating between COVID-19 convalescent and negative control samples.**

| Peptide Sequence | Protein | Start Position | End Position | Reactive Convalescent Samples |
|---|---|---|---|---|
| SFKEELDKYFKNHTSPDVDLGDISGINASV | S | 1147 | 1176 | 12 |
| SKPSKRSFIEDLLFNKVTLADAGFIKQYGD | S | 810 | 839 | 9 |
| NAAIVLQLPQGTTLPKGFYAEGSRGGSQAS | N | 154 | 183 | 8 |
| GDAALALLLLDRLNQLESKMSGKGQQQQGQ | N | 215 | 244 | 3 |

**Table S3. Acknowledgement of GISAID sequences used in the design of SCV2 PepSeq Library.**

Available at github.com/jtladner/Manuscripts/tree/master/2020_Ladner_SARS2PepSeq.
